# *De novo* annotation and characterization of the translatome with ribosome profiling data

**DOI:** 10.1101/137216

**Authors:** Zhengtao Xiao, Rongyao Huang, Yuling Chen, Haiteng Deng, Xuerui Yang

**Author notes:** Corresponding author: X.Y.

## Abstract

By capturing and sequencing the RNA fragments protected by translating ribosomes, ribosome profiling sketches the landscape of translation at subcodon resolution. We developed a new method, RiboCode, which uses ribosome profiling data to assess the translation of each RNA transcript genome-wide. As shown by multiple tests with simulated data and cell type-specific QTI-seq and mass spectrometry data, RiboCode exhibits superior efficiency, sensitivity, and accuracy for *de novo* annotation of the translatome, which covers various types of novel ORFs in the previously annotated coding and non-coding regions and overlapping ORFs. Finally, to showcase its application, we applied RiboCode on a published ribosome profiling dataset and assembled the context-dependent translatomes of yeast under normal condition, heat shock, and oxidative stress. Comparisons among these translatomes revealed stress-activated novel upstream and downstream ORFs, some of which are associated with potential translational dysregulations of the main protein coding ORFs in response to the stress signals.

## Introduction

Ribosome profiling, also called Ribo-seq, generates genome-wide maps and quantifications of the ribosome protected RNA fragments (RPF)(Ingolia et al., 2009), which provide real-time snapshots of translation (translatome) across the whole transcriptome. Many studies have exploited this powerful technique to characterize the landscapes of translation, including the translational rates(Hsieh et al., 2012; Su et al., 2015; Thoreen et al., 2012), pausing upon stress signals(Gerashchenko et al., 2012; Liu et al., 2013; Shalgi et al., 2013), stop codon read-through (Dunn et al., 2013), translation potential of non-coding sequences(Bazzini et al., 2014; Fritsch et al., 2012; Ingolia et al., 2011; Ji et al., 2015), and alternative reading frames(Fritsch et al., 2012; Menschaert et al., 2013). However, it has also been frequently shown that the ribosome occupancy itself, as indicated by the RPF reads mapped on the transcriptome, is not sufficient for the calling of the active translation, given the possibilities of data and experimental noise, regulatory RNAs that bind with the ribosome, and ribosome engagement without translation(Banfai et al., 2012; Guttman et al., 2013). This therefore requires a specially designed methodology for processing the ribosome profiling data to recover the active translation events from the usually distorted and ambiguous signals. Such method should fully account for the complexity of translation itself, such as alternative initiation sites and overlapping open reading frames (ORFs).

Owing to its subcodon resolution, ribosome profiling reveals the precise locations of the peptidyl-site (P-site) of the 80S ribosome in the RPF reads. Aligned by their P-site positions, the RPF reads resulting from the translating ribosomes should therefore exhibit 3-nt periodicity along the ORF, which is the strongest evidence of active translation. Only recently have different strategies been developed to assess the translation by testing this 3-nt periodicity of ribosome engagement(Bazzini et al., 2014; Calviello et al., 2016; Duncan and Mata, 2014; Michel et al., 2012), and only a couple of these methods were designed for *de novo* translatome annotation(Bazzini et al., 2014; Calviello et al., 2016). In general, and as shown later in the present study, various limitations hinder the broad applications of these methods. Therefore, we have developed a new statistically vigorous method, RiboCode, for the *de novo* annotation of the full translatome by quantitatively assessing the 3-nt periodicity (Fig. 1). Tested using both simulated and real data, RiboCode exhibited superior efficiency, sensitivity, and accuracy to the existing methods. In addition, unlike other methods, the methodology design of RiboCode allows the assessment of non-canonical translations such as overlapping ORFs. Finally, to showcase the application of RiboCode in reconstructing the context-dependent translatomes, we applied RiboCode on a published ribosome profiling dataset to assemble the translatomes of yeast under normal and stress conditions(Gerashchenko and Gladyshev, 2014). Comparisons among these translatomes revealed novel ORFs in the canonically non-coding regions that were activated in response to heat shock and oxidative stress. Quantitative analysis of the ORFs further showed that some of the upstream ORFs (uORFs) and downstream ORFs (dORFs) were indeed associative with the potential translation dysregulation of the coding regions of the same mRNA transcripts.

**Figure 1.**
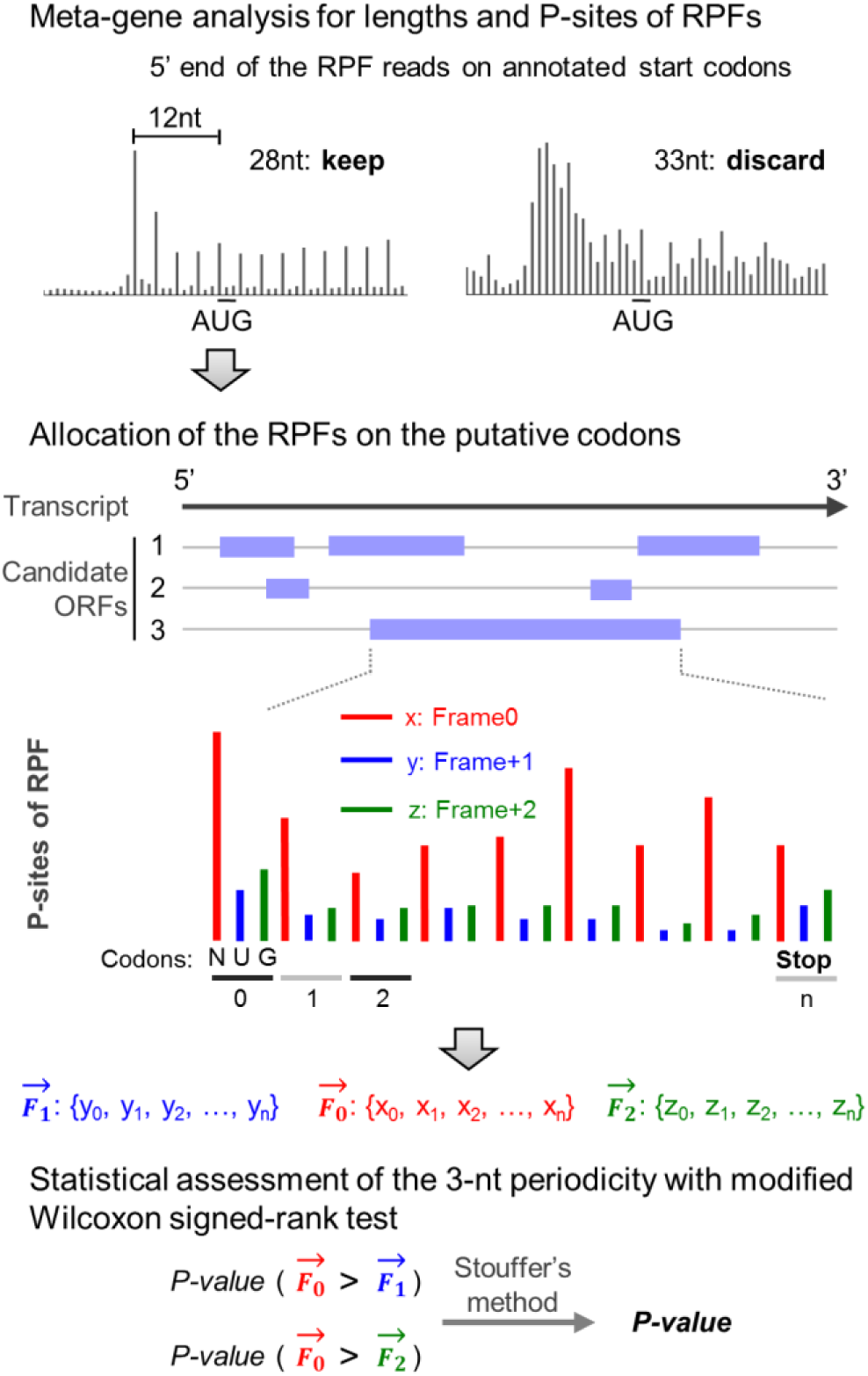
The methodology design of RiboCode. Schematic description of RiboCode. Further details are provided in the Methods section.

## Results and Discussions

### Methodology design of RiboCode

The methodology of RiboCode primarily relies on evaluation of the 3-nt periodicity of the RPF reads aligned by the P-sites on the RNA sequence that is being actively translated. Considering the usually distorted patterns of RPF read mapping and potentially high noise level of the ribosome profiling data, we adapted the Wilcoxon signed-rank test to assess the oddness of consistently higher in-frame reads along the whole ORF. We chose this strategy because of the following reasons. First, this analysis is insensitive to the potentially strong artificial in-or off-frame RPF signals at small fractions of the codons in the whole ORF. Such artifacts are not rare in ribosome profiling data, due to the occasional RPF read duplicates potentially resulted from the PCR amplification bias. Second, the RPF reads are often sparse along the ORF, and this test is not distracted by the codons with no RPF read, i.e., no evidence for either active translation or the opposite. Third, this test is insensitive to background noise due to RNA contamination or incorrect P-site allocation. Last but not the least, this statistical test is computationally cost-effective, thereby rendering high computation efficiency of RiboCode.

The workflow of RiboCode is composed of 3 major steps, 1) preparing the transcriptome for search of the candidate ORFs, 2) determining the length range of the RPF reads that are most likely to have resulted from active translation, and identifying the P-site positions in these reads, and 3) assessing the active translation event via statistical comparisons among the 3 vectors representing the RPF read densities in and off the reading frame along each candidate ORF. The analysis strategy of RiboCode is illustrated in Fig. 1, and the details of the method design are provided in the methods section.

### Benchmark the performance of RiboCode

To benchmark the performance of RiboCode and the existing methods designed for *de novo* annotation of the translatome, including RiboTaper(Calviello et al., 2016) and ORFscore(Bazzini et al., 2014), we performed test runs with a published ribosome profiling dataset in human HEK293 cell(Gao et al., 2015). RPF reads of the consensus coding sequence (CCDS) exons were used as positives, and the paralleled RNA-seq data was included to mimic the negatives, i.e., reads of untranslated RNA that lack the 3-nt periodicity. We calibrated the cutoffs for all three methods, RiboCode, RiboTaper, and ORFscore, to achieve the same false positive rate (∼7.5%, 3215). As a result, RiboCode recovered many more CCDS exons than did the other two methods (Fig. 2a, detailed results in Supplementary Table 1).

**Figure 2.**
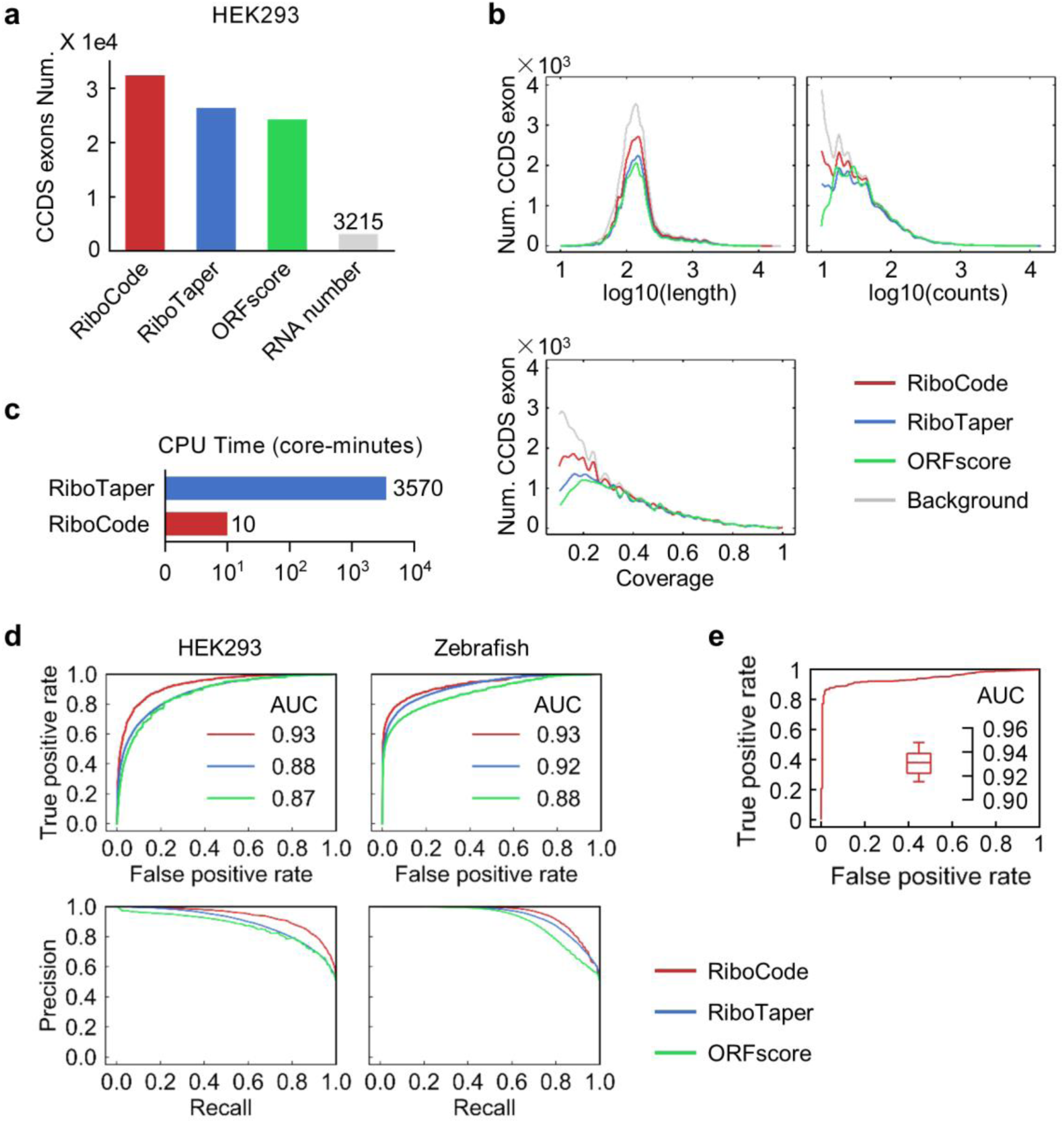
General performance of RiboCode benchmarked with simulation data. (a) The numbers of CCDS exons identified by different methods with the RPF data in HEK293 cells. The cutoffs used for three methods, RiboCode, RiboTaper, and ORFscore, were calibrated so that they produced the same numbers of false positives with the RNA-seq data. (b) Distributions of the lengths, total read counts, and coverage of the CCDS exons identified by RiboCode, RiboTaper, and ORFscore, based on the same false positive rate. (c) The CPU time (in core-minutes) consumed for *de novo* annotation of the translatome from a human ribosome profiling dataset, with RiboTaper and RiboCode. (d) ROC and precision curves generated with the results of RiboCode, RiboTaper and ORFscore with two simulation datasets, one generated from the HEK293 cell data in Gao et al (left) and the other one from the Zebrafish data in Bazzini et al (right). (e) A representative ROC curve generated with the results of RiboCode on a simulation data set specifically for the overlapping ORFs. The box plot inside summarizes the AUC of 20 ROC curves from 20 such simulation data sets processed by RiboCode.

In addition, unlike the other methods, RiboCode yielded significant distinctiveness when processing the RPF reads and RNA-seq reads, which is not or only slightly dependent on the length, read counts, or coverage of the CCDS exons (Supplementary Fig. 1a-c). Indeed, the distributions of the lengths, RPF read counts, and coverages of the results are highly concordant with those of the full CCDS set as a background (Fig. 2b), which suggests that RiboCode is capable of comprehensively annotating the full translatome. Note that the exons with the RPF read count fewer than 10 or the coverage smaller than 0.1 were discarded. The other two methods, however, showed some bias towards the ORFs with high read counts and coverage (Fig. 2b, Supplementary Fig. 1a-c).

Finally, it is worth noting that owing to the efficient statistical design, RiboCode is very user-friendly and requires little computation resource. Annotation of the full translatome with the ribosome profiling dataset in HEK293 cells(Gao et al., 2015) took about 10 minutes with RiboCode on a single-core computer (10 core-minutes), which is trivial compared to the ∼20 hours on a 16-core high-performance computing platform (3570 core-minutes) – 99.7% reduction of the computation burden (Fig. 2c).

The tolerance to the sometimes unavoidable technical noise is important for the potential applications of a method. This is especially true for the analysis of ribosome profiling data, given its nature of high noise resulting from contaminations of non-ribosome-bound RNA, regulatory RNA in the ribosomal complex, inappropriate RPF read length selections, and inaccurate P-site position. These noises result in either contamination of the RPF reads or incorrect alignments of the reads, both of which should weaken the 3-nt periodicity. Essentially, this can be simulated by shuffling the P-site among the 3 positions, -1, 0, or +1 in relative to the original position, for a randomly selected subset of the RPF reads, which by definition destroys the 3-nt periodicity of this RPF subset. Stress tests of the three methods applied on such datasets generated from the HEK293 data(Gao et al., 2015), in which different percentages of the RPF reads were disturbed, showed that RiboCode consistently out-performed the other two methods with low-to high-noise data (Supplementary Fig. 2a). In addition, considering that the sequencing depth of the different published ribosome profiling studies could vary significantly, we also tested the performance of the three methods with different numbers of RPF reads. As Supplementary Fig. 2b shows, RiboCode was able to deliver relatively good sensitivity and specificity, which were not much sacrificed with fewer total RPF reads. Taken together, these tests suggest that RiboCode is of great value for annotating the translatomes with published ribosome profiling datasets that are of low quality or with limited number of usable RPF reads.

### Comparisons between RiboCode and existing methods

To further systematically evaluate the sensitivity and specificity of the three methods, we prepared ROC curves with the results of the different methods applied on two published ribosome profiling datasets(Bazzini et al., 2014; Gao et al., 2015), one in HEK293 cells and the other in Zebrafish (Fig. 2d). The paralleled RNA-seq data was again used as true negatives. The detailed results are provided in Supplementary Tables 1. These test runs illustrated the superior sensitivity and specificity of RiboCode compared to the two other existing methods (Fig. 2d). The P-value distributions of the results of RiboCode indeed showed a much cleaner separation of the CCDS exons called from the RPF reads and the ones from the RNA-seq reads (Supplementary Fig. 1d).

One of the major challenges for the *de novo* annotation of the translatome is the complicated re-coding events, including the frequently found overlapping off-frame ORFs. The methodology design of RiboCode genuinely allows assessment of the overlapping ORFs, while the two existing methods for *de novo* translatome annotation, RiboTaper and ORFscore, have difficulties in recovering such recoding events. Here, we used a simulation dataset to test the performance of RiboCode in annotating the actively translated overlapping ORFs. Specifically, with the previously used HEK293 dataset(Gao et al., 2015),we overlaid the RPF reads of two annotated CCDS with a +1 or +2 frame shift to simulate the RPF reads from an artificial pair of overlapping ORFs. For a negative case, we randomly assigned an artificial ORF that has a frame shift and overlaps with an annotated CCDS. As Fig. 2e shows, RiboCode exhibited high sensitivity and accuracy in capturing the actively translated overlapping ORFs. RiboCode is therefore the first method for *de novo* annotation of the translatome including overlapping ORFs, which is the major form of recoding.

### Validations of the predicted ORFs by QTI-seq data

Multiple studies have reported widespread alternative translation initiation(Ingolia et al., 2011; Kozak, 1989; Lee et al., 2012), which is suspected to be context-dependent. A precise annotation of the translation initiation sites is therefore critical for the complete assembly of the translatome. Experimentally, blockage of elongation from the newly assembled initiation complex with antibiotics such as harringtonine and lactimidomycin(Ingolia et al., 2011; Lee et al., 2012) allows the efficient screening of the translation initiation sites with ribosome profiling. However, such an experimental setting is not applicable in most of the previous and, potentially, future studies. Therefore, it would be greatly beneficial to have a method that can precisely allocate at least some of the translation initiation sites from the regular ribosome profiling data.

We used a QTI-seq dataset that comprehensively mapped the translation initiation sites in HEK293 cells(Gao et al., 2015) to test the performance of RiboCode and RiboTaper in correctly annotating the real start codons with the ribosome profiling data in the same cellular context. The results of RiboCode and RiboTaper, for a complete *de novo* translatome annotation with the regular ribosome profiling data in HEK293 cells(Gao et al., 2015), were provided in Supplementary Table 2, in which the detailed information of the ORFs including the initiation sites can be found. The accumulation curves were prepared, thus indicating the proportions of the presumably true initiation sites (identified by QTI-seq) that were correctly recovered by the two methods with the ribosome profiling data (Fig. 3a). It appears that RiboCode is slightly more efficient in annotating the translation initiation sites. For example, from the same number of identified ORFs, a Venn diagram showed a larger overlap between the initiation sites of the ORFs identified by RiboCode and the initiation sites from QTI-seq (Fig. 3a).

**Figure 3.**
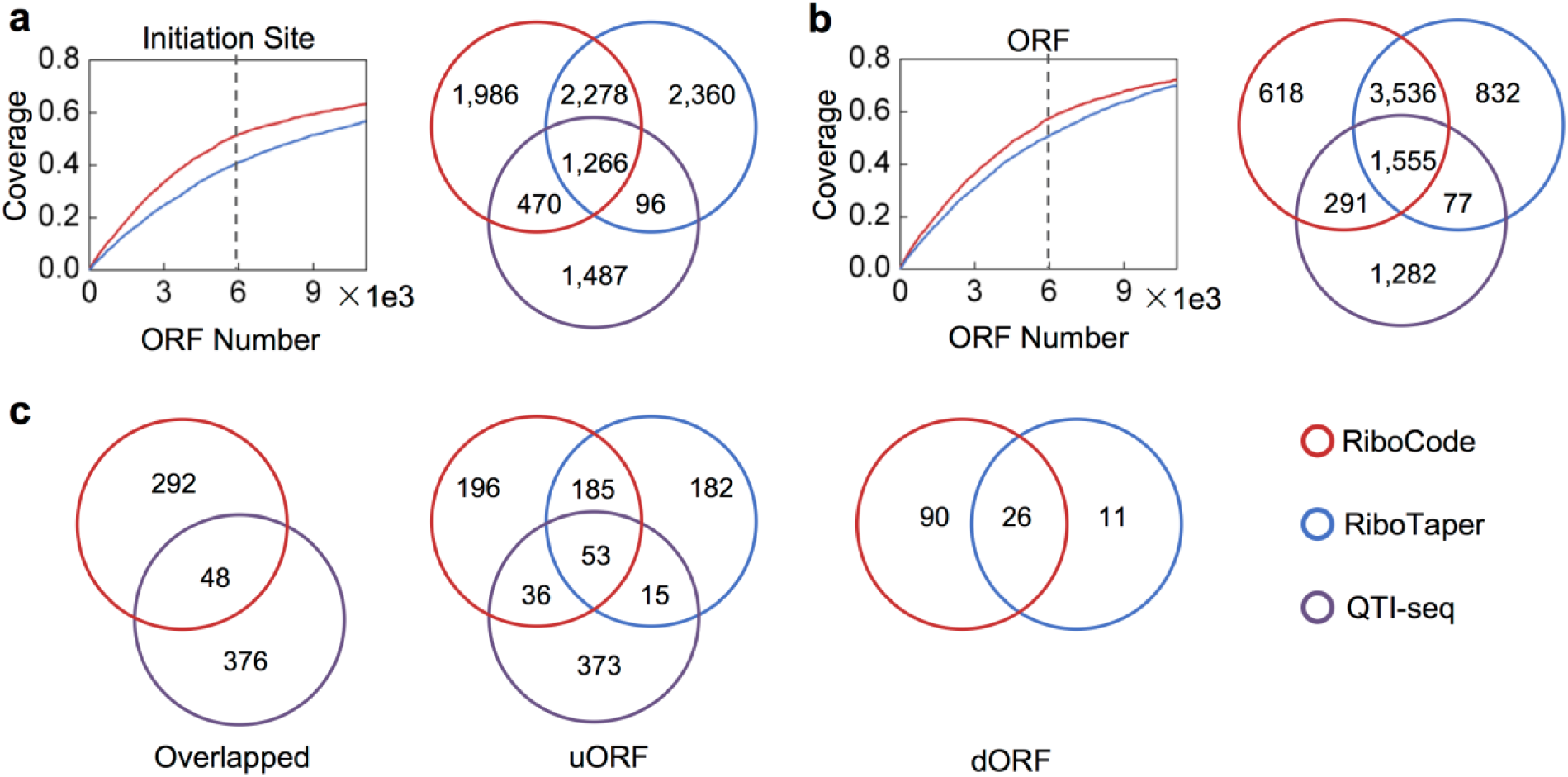
Validation of the ORFs with QTI-seq data. (a, b) Accumulation curves showing proportions of the initiation sites (a) and the annotated ORFs (b) identified by QTI-seq that were recovered by RiboCode or RiboTaper. The Venn diagrams to the right show the overlaps among the 3 sets of initiation sites (a) or annotated ORFs (b), one identified from QTI-seq and two from ribosome profiling data processed by RiboCode and RiboTaper. The P-value cutoffs were selected for RiboCode and RiboTaper so that they yielded the same total number of positive hits as marked on the accumulation curves. (c) Venn diagrams showing the overlaps among the uncanonical ORFs (overlapping ORFs, uORFs, and dORFs) identified from the QTI-seq data and from the ribosome profiling data processed by RiboCode and RiboTaper.

By capturing the accumulated ribosomes at the initiation sites due to stalled translation elongation, the QTI-seq data was also used to predict the actively translated ORFs, including both the annotated coding sequence (CDS) and unannotated ORFs, such as the upstream ORFs (uORFs) and the previously discussed overlapping ORFs. We then evaluated the overlaps among the ORFs identified by RiboCode and RiboTaper with ribosome profiling data and the ORFs inferred from the QTI-seq data. First, for the previously annotated ORFs from the CDS regions, the accumulation curves, which indicate the proportion of the ORFs from the QTI-seq data that were also supported by RiboCode or RiboTaper with the ribosome profiling data, illustrated the higher sensitivity of RiboCode in recovering the annotated ORFs identified by QTI-seq (Fig. 3b). Again, with the same total number of ORFs being reported, a Venn diagram showed that with the ribosome profiling data in the same cell context, RiboCode yielded larger overlaps than RiboTaper did, with the ORFs inferred from the QTI-seq data (Fig. 3b). Finally, Venn diagrams specifically for the overlapping ORFs, the uORFs, and the downstream ORFs (dORFs) were also prepared (Fig. 3c). As discussed previously, RiboTaper did not identify any overlapping ORFs (Fig. 3c).

### The translatomes assembled by RiboCode and support from MS data

Taken together, the results above illustrate the sensitivity and accuracy of RiboCode for comprehensive *de novo* annotation of the translatome with ribosome profiling data. We then summarized the different types of ORFs recovered by RiboCode and RiboTaper, with two published ribosome profiling datasets in the HEK293 cell(Gao et al., 2015) and Zebrafish(Bazzini et al., 2014) (Fig. 4a, b). The detailed results are provided in Supplementary Tables 2 (HEK293) and 3 (Zebrafish). The protein or peptide products from these ORFs were further validated, in a cell type-specific manner, with published Mass Spectrometry (MS) data of the HEK293 cell and Zebrafish (Fig. 4c, d, Supplementary Table 4). Under various categories of the ORFs, large proportions of the translatomes assembled by RiboCode, but much smaller by RiboTaper, were supported by the MS data (Fig. 4c, d). These validated ORFs include many previously unannotated uORFs, dORFs, overlapping ORFs, novel ORFs from annotated coding and non-coding genes, many of which were only identified by RiboCode, but not by RiboTaper. Examples were given in Supplementary Fig. 3a-e.

**Figure 4.**
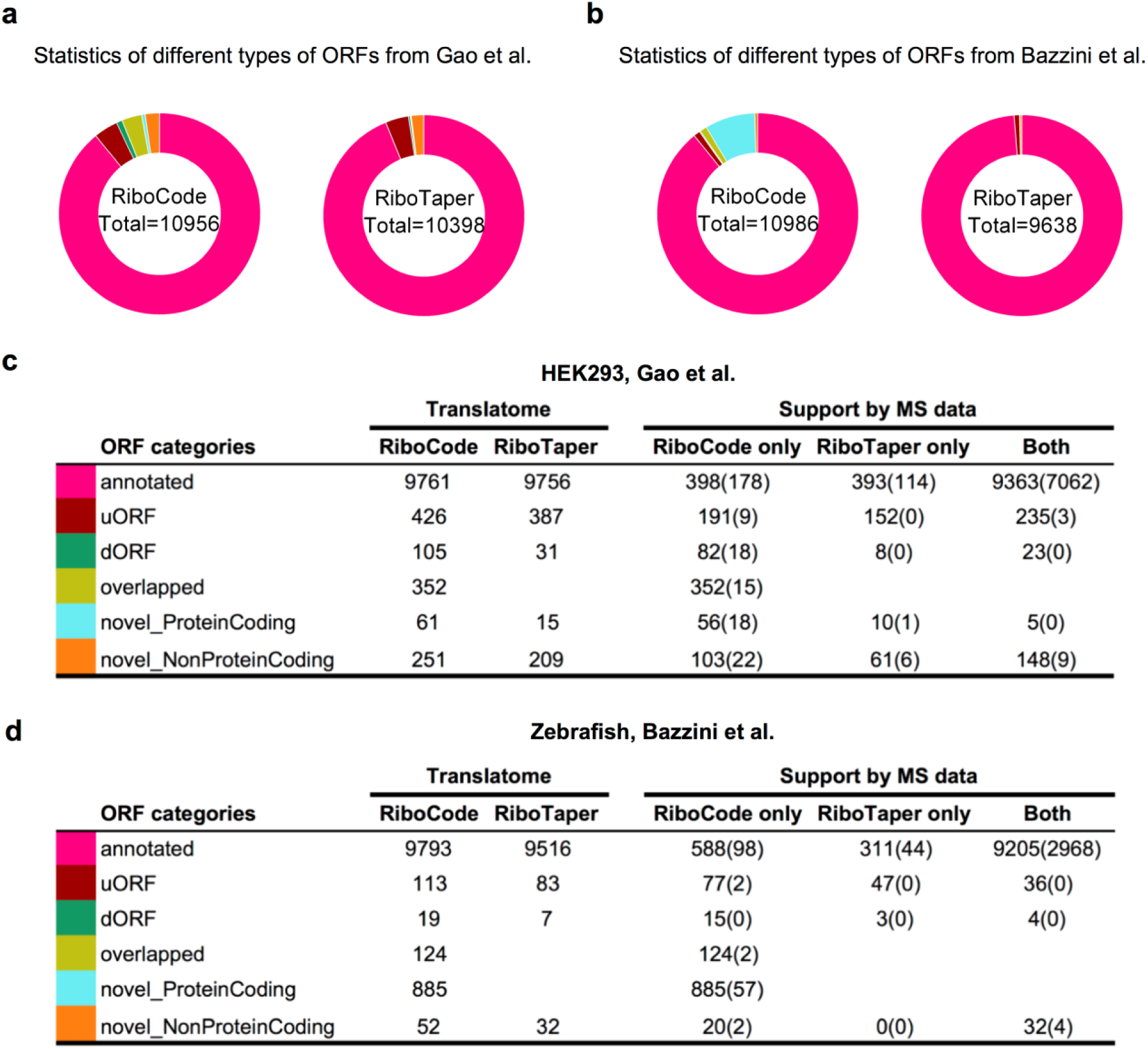
*De novo* annotations of the full translatomes and validations with MS data. (a, b) The composition of the translatomes assembled by RiboCode and RiboTaper, with the HEK293 dataset (a) and the Zebrafish dataset (b). (c, d) Statistics of the ORFs in different subcategories identified by RiboCode and RiboTaper, in HEK293 cell (c) and Zebrafish (d). Numbers of the ORFs, identified by RiboCode only, RiboTaper only, and by both of them, which have support from the MS data, were given in the parentheses.

### Application of RiboCode in assembling the context-dependent translatomes of yeast

We used the ribosome profiling data in yeast under 3 conditions: normal, heat shock, and oxidative stress, to showcase the application of RiboCode for *de novo* assembly of the context-specific translatomes. Fig. 5a, b summarized the yeast translatomes under the 3 conditions (details of the annotated ORFs are provided in Supplementary Table 5). In general, RiboCode identified more uORFs and dORFs being translated upon heat shock and oxidative stress. Next, we compared the RPF read counts of the different ORF types in the translatomes between the stress and normal conditions. The raw and normalized RPF read counts of all the ORFs annotated by RiboCode are provided in Supplementary Table 6. While the previously annotated protein coding genes have similar overall distributions of the RPF read counts, the uORFs and dORFs showed markedly higher RPF read counts under the stress conditions (Fig. 5c, Supplementary Fig. 5). This is well in line with previous knowledge about translation of uORFs in multiple organisms in response to various stress signals(Brar et al., 2012; Gerashchenko et al., 2012; Laing et al., 2015; Starck et al., 2016; Young and Wek, 2016).

**Figure 5.**
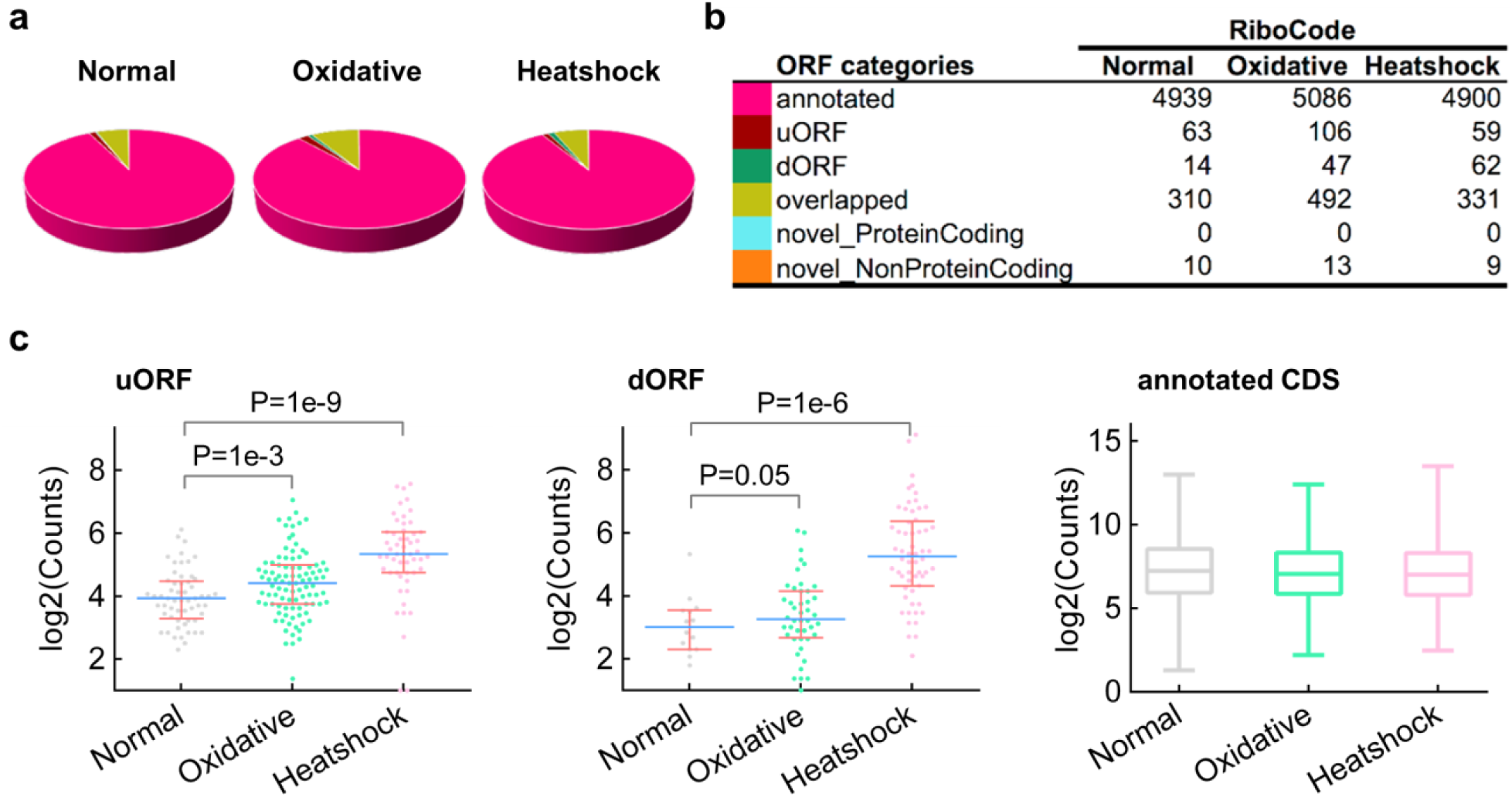
Application of RiboCode for assembly of the yeast translatomes under normal and stress conditions. (a, b) The composition of the translatomes assembled by RiboCode, with the ribosome profiling data of yeast, under normal condition, oxidative stress, and heat shock. (c) Distributions of the normalized RPF read counts (log2) of the uORFs, dORFs, and annotated CDS, under the three conditions: normal, oxidative stress, and heat shock. Mann–Whitney U tests were performed to assess the statistical significance of the difference between the distributions of uORF or dORF under heat shock vs. normal or oxidative stress vs. normal condition. The P-values were provided in the figure.

Next, we looked into the RPF read counts of the main coding ORFs that are at downstream of the uORFs or upstream of the dORFs. Fig. 6a-d showed the fold changes of these main coding ORFs upon the stress signals, on the background of all the annotated CDS regions. The vertical bars, representing each of these main coding ORF, were color-coded based on the fold change of its uORF (Fig. 6a, b) or dORF (Fig. 6c, d) under the heat shock (Fig. 6a, c) or oxidative stress (Fig. 6b, d) condition, compared to the normal condition. It appears that the translational up-regulation of some uORFs or dORFs were associated with potential stress-induced translational repression of the annotated coding ORF of the same transcripts (see the red vertical bars to the left side of the spectrums in Fig. 6a-d, and some examples shown in Fig. 6e-j). Indeed, many previous studies have reported that activation of the uORFs could result in translational inhibition of the downstream coding ORF(Brar et al., 2012; Gerashchenko et al., 2012; Ingolia et al., 2011; Lee et al., 2012). On the other hand, many of the uORFs or dORFs were positively associated with the translation of the main coding ORF (see the red vertical bars to the right side and the blue bars to the left side of the spectrums in Fig. 6a-d). This could be attributed to the general translational or transcriptional up-regulation of the mRNA transcripts that harbor the main protein-coding ORF and the uORF or the dORF. More data and further analysis would be needed to fully elucidate the potential involvements of the uORFs and dORFs in regulating the translation of the main protein coding ORFs.

**Figure 6.**
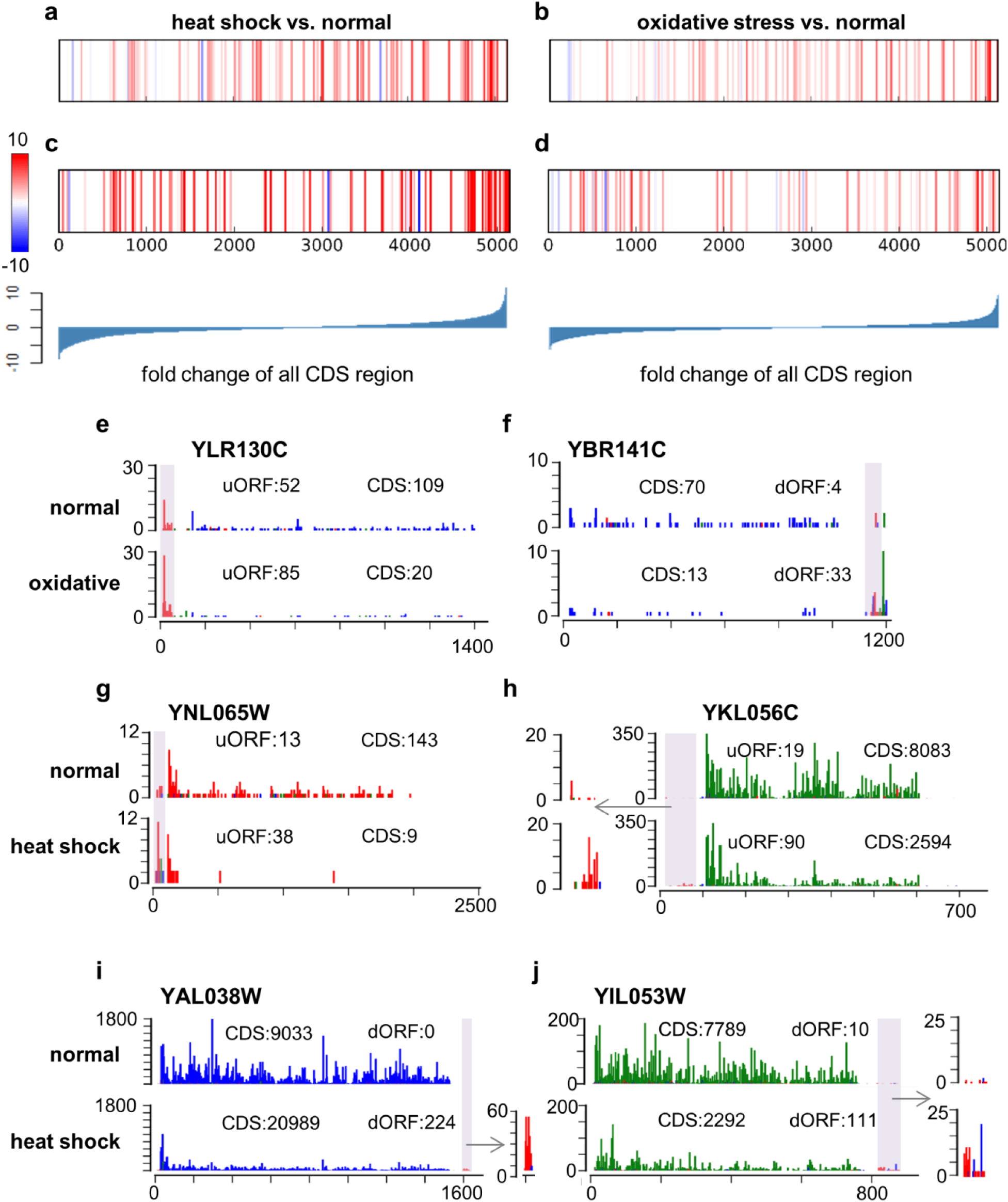
Associations of the uORFs and dORFs with the main protein coding ORF. (a-d) All the annotated canonical protein coding ORFs were sorted based on the fold change of their normalized RPF read counts under the stress condition (a and c for oxidative stress, and b and d for heat shock) vs. the normal condition. On this background, the ORFs with upstream uORFs (a, b) or downstream dORFs (c, d) in the same mRNA transcripts were marked as vertical bars. The color of these bars represents the fold change of the RPF read counts on the uORF (a, b) or dORF (c, d). (e-j) 6 examples of the uORF (e, g, h) or dORF (f, i, j) that appear negatively associated with the main ORF in response to oxidative stress (e, f) or heat shock (g-j). The colored bar on each nucleotide position represents the count of RPF reads allocated according to its P-site position. Three different colors represent the three frames. The total RPF read counts of the main protein coding ORFs and the uORFs or dORFs are given in the figures.

## Conclusions

The collection of ribosome profiling data has been quickly increasing, thus shaping the landscape of translation in various systems with increasing detail. There is a clear need for *de novo* annotations of the species- and cellular context-dependent translatomes, which have largely lagged behind the genome and transcriptome annotations. Designed for the comprehensive *de novo* annotation of the translatome with ribosome profiling data, RiboCode presents remarkable advantages over its counterparts. It has significantly higher efficiency and accuracy in calling the actively translated ORFs. Its capability of recovering recoding events such as overlapping ORFs and its consistent performance, which is largely independent of the length, read count and coverage, assure the comprehensiveness of the translatome annotation. In addition, RiboCode’s relatively consistent performance with different noise levels and sequencing depths is another valuable feature for processing the various published ribosome profiling data. Last but not least, RiboCode requires very little computational resource, thereby enabling routine large-scale translatome annotation with ribosome profiling datasets. Therefore, we are presenting RiboCode to the community for the processing of published and future ribosome profiling data to obtain more comprehensive understanding of the context-specific translatomes.

## Methods

### Ribosome profiling and RNA-seq data preprocessing

The two sets of ribosome profiling data, one in HEK293 cell(Gao et al., 2015) and the other in Zebrafish(Bazzini et al., 2014), were downloaded from the NCBI Sequence Read Archive for HEK293 (accession ID SRA160745) and the Gene Expression Omnibus (GEO) database for Zebrafish (accession ID GSE53693). The ribosome profiling data of yeast under normal, oxidative stress, and heat shock conditions was downloaded from GEO (GSE59573).

The pre-processing procedure of the ribosome profiling data has been described previously (Xiao et al., 2016). Specifically, the cutadapt program (Martin, 2011) was used to trim the 3’ adaptor in the raw reads of both mRNA and RPF. Low-quality reads with Phred quality scores lower than 20 (>50% of bases) were removed using the fastx quality filter (http://hannonlab.cshl.edu/fastx_toolkit/). Next, sequencing reads originating from rRNAs were identified and discarded by aligning the reads to rRNA sequences of the particular species using Bowtie (version 1.1.2) with no mismatch allowed. The remaining reads were then mapped to the genome and spliced transcripts using STAR with the following parameters: --outFilterType BySJout --outFilterMismatchNmax 2 --outSAMtype BAM --quantMode TranscriptomeSAM --outFilterMultimapNmax 1 --outFilterMatchNmin 16. To control the noise from multiple alignments, reads mapped to multiple genomic positions were discarded.

For the yeast data, the RPF reads on each ORF was counted with custom scripts based on HTSeq-count (Anders et al., 2015; Xiao et al., 2016) in intersection-strict mode. Only the RPF reads with length between 27 and 29 nt, which were found to exhibit strong 3-nt periodicity, were used for counting. Due to the potential accumulation of ribosomes around the starts and ends of coding regions (Ingolia et al., 2014; Ingolia et al., 2011), reads aligned to the first 15 and last 5 codons were excluded for counting of RPF reads for the ORFs longer than 100 nt. The raw read counts of each ORF across the three conditions were further subjected to median-of-ratios normalization (Anders et al., 2012).

### RiboCode step 1: preparation of the transcriptome annotation

This step defines the annotated transcripts, from which the candidate ORFs will be identified. This is done by the *prepare_transcripts* command in the RiboCode package, with inputs of a GTF file and a genome FASTA file. As examples, in the present study, the GTF and FASTA file (release 74 for human, and release 87 for Zebrafish) were downloaded from the Ensembl FTP repository (http://www.ensembl.org/info/data/ftp/index.html). Each transcript was assembled by merging the exons according to the structures defined in the GTF file. The transcript sequences were then retrieved from the genome FASTA file.

### RiboCode step 2: filtering of the RPF reads and identification of the P-site locations

The purpose of this step, with the *metaplots* command in the RiboCode package, is to 1) select the length range of the RPF reads that most likely originated from the translating ribosomes and 2) identify the P-site locations for different lengths of the RPFs. This was done with a meta-gene analysis of the RPF reads mapped on the previously annotated coding genes. Specifically, for each set of the RPF reads with a particular length, the distances from their 5’ ends to the annotated start and stop codons were calculated and summarized as histograms (Supplementary Fig. 4 as an example). The length range, in which the pooled RPF reads showed strong 3-nt periodicity from their 5’ ends to the start and stop codons, should then be determined by the user, for the following analysis of RiboCode. In the examples shown in Supplementary Fig. 4, the RPF length range was deemed to be 26-29 nt for HEK293 and 28-29 nt for Zebrafish.

Also from the histograms for each of the RPF lengths selected above, the P-site locations were inferred according to the offsets of the 5’ end of the RPF reads mapped on the start codons. In the examples shown in Supplementary Fig. 4, the P-sites were identified as the +12^th^ nt of all the RPF reads within the selected length range for HEK293 and Zebrafish data. Supplementary Table 7 presented the selected read lengths and the P-site positions for the different ribosome profiling datasets used in the present study.

Based on our experience, in most cases, selection of the RPF reads around 28-30 nt is generally appropriate, and their P-site positions are usually at +12. However, we believe that it is critical to run this step of RiboCode to extract the RPFs that are most likely from the translating ribosomes and to precisely determine their P-site positions. Alternatively, the users have the option to skip this step and directly provide the information of read length and P-site positions based on their experiences, although this is not recommended, especially when the experimental conditions (species, culturing condition, stress) or the procedure of ribosome profiling (nuclease, buffer, library preparation) have been changed.

### RiboCode step 3: identification of the candidate ORFs and assessment of the 3-nt periodicity

As the primary analysis procedure of RiboCode, this step is executed with a single command *RiboCode*. It starts with a transcriptome-wide search for the candidate ORFs that start from a canonical start codon (AUG) and end at the next stop codon. Optionally, alternative start codons provided by the users, for example CUG and GUG, can also be included in the search for the candidate ORFs in the regions outside of the ORFs with the canonical start codon AUG. Next, based on the mapping results of the RPF reads within the length range identified in the second step, for each nucleotide of the candidate ORF, RiboCode counts the number of reads, of which the P-sites were allocated on the particular nucleotide. Eventually, RiboCode generates a spectrum of the P-site densities at each nucleotide along each candidate ORF.

Mathematically, the spectrum of the P-site densities along each candidate ORF is a numerical vector with the length of the ORF. From this vector, we simply derived 3 shorter vectors, each with 1/3 of the length of the ORF. As shown in Fig. 1, one of these 3 vectors, *F*_*0*_, represents the P-site density along the first nucleotide of each codon, from the start to the stop codon. Similarly, the other two vectors, *F*_*1*_ and *F*_*2*_, represent the P-site densities along the 2^nd^ and the 3^rd^ nucleotide, respectively, of each codon. To assess the 3-nt periodicity, the Wilcoxon signed rank test strategy was modified and used to evaluate whether *F*_*0*_ is generally greater than *F*_*1*_ and *F*_*2*_ at the non-zero positions. Accordingly, this would yield two *P-values*, indicating the significance levels of *F*_*0*_ *> F*_*1*_ and *F*_*0*_ *> F*_*2*_. Finally, an integrated P-value was derived with Stoufer’s method, which represents the overall statistical significance of the 3-nt periodicity.

Most CDS have multiple start codons upstream of the stop codon, and we followed two simple principles to identify the translation initiation sites for the candidate ORFs. (1) We used the same procedure of the modified Wilcoxon signed rank test, as described above, to assess the 3-nt periodicity of the RPF reads mapped between the most upstream (first) start codon and the next one (second) downstream. This was done only if there were more than 10 codons in this region, of which the in-frame RPF counts are larger than zero. If this test resulted in a statistically significant 3-nt periodicity (P-value smaller than the cutoff provided by the user, for example 0.05), we defined the first start codon as the translation initiation site. Otherwise, we disregard it and start the same procedure over for the region between the second start codon and the subsequent one. (2) If there are limited RPF reads (fewer than 10 codons with none-zero in-frame RPF counts) between two neighboring start codons, we chose the upstream start codon of the region, in which the codons that have more in-frame than off-frame RPF reads (frame0 > frame 1 and frame0 > frame2) are greater than the ones that do not (frame0 <= frame 1 and frame0 <= frame2).

### Preparation of the simulation data

The exon-level simulation datasets used in Fig. 2a, b, d were generated from the two datasets of ribosome profiling with RNA-seq in parallel, in HEK293 and Zebrafish. The P-site data track for each CCDS exon from the Ensembl annotation was created using the RiboTaper package (P_sites_all_tracks_ccds and Centered_RNA_tracks_ccds files in data_tracks generated by RiboTaper). The read lengths and P-site locations used in these data are provided in Supplementary Table 7. For the RNA-seq data used as true negatives, the 25^th^ position was arbitrarily defined as the P-site position. Exons shorter than 10 nt were discarded. The RiboTaper package was used to calculate the ORFscore and *P-value* of RiboTaper (results_ccds generated by RiboTaper).

The ribosome profiling datasets with different levels of noise, used in Supplementary Fig. 2a, were generated by subsampling different fractions of the RPF reads of the HEK293 data(Gao et al., 2015) and shuffling their P-site positions among -1, 0, and +1 in relative to the original position (+12 nt). For the datasets with reduced sequencing depth, used in Supplementary Fig. 2b, we just randomly discarded different percentages of the RPF reads in the HEK293 data(Gao et al., 2015).

### Annotation of the ORFs from QTI-seq data

For each initiation site identified by the QTI-seq data(Gao et al., 2015), we selected the closest downstream in-frame stop codon, thereby annotating an ORF. If one initiation site has more than one in-frame stop codon in different transcripts of the same gene, only the one harbored in the longest transcript was chosen.

### Mass spectrometry data collection and processing

Human MS/MS data of HEK293 cells were obtained from our previously published study(Wang et al., 2016) and ProteomeXchange Consortium(PXD002389). Zebrafish MS/MS data was downloaded from ProteomeXchange Consortium (PXD000479, tissue of Testis). The peptides were searched using the SEQUEST searching engine of Proteome Discoverer (PD) software (version 1.4). The same search criteria as published before(Wang et al., 2016) was used. The false discovery rate (FDR), calculated using Percolator provided in PD, was set to 0.1 for peptides and proteins.

### Software availability

The RiboCode package is available at https://pypi.python.org/pypi/RiboCode. A detailed step-by-step instruction of the data preprocessing and usage of RiboCode is also provided. The method requires a genome FASTA file, a GTF file for transcriptome annotation, and the alignment result file of the ribosome profiling data.

## Acknowledgments

The authors wish to acknowledge the support from the Gene Sequencing, Protein Chemistry, and Computing core facilities at the National Protein Science Facility (Beijing) and the Center for Biomedical Analysis of Tsinghua University. This work was supported by the National key research and development program, Precision Medicine Project (2016YFC0906001 to X.Y.), the National Natural Science Foundation of China (91540109 and 81472855 to X.Y.), the Tsinghua University Initiative Scientific Research Program (20131089278 to X.Y.), the Tsinghua–Peking Joint Center for Life Sciences, and the 1000 talent program (Youth Category).

### Authors’ Contributions

Z.X. and X.Y. conceived and designed the study. Z.X. developed the algorithm and performed the analyses with help from R.H. Y.C. and H.D. performed the mass spectrometry data analysis. X.Y. supervised the whole project. Z.X. and X.Y. wrote the manuscript. All authors have read and approved the final manuscript.

### Competing interests

The authors declare no financial conflict of interest.

## Supplementary Figures

**Supplementary Figure 1.**
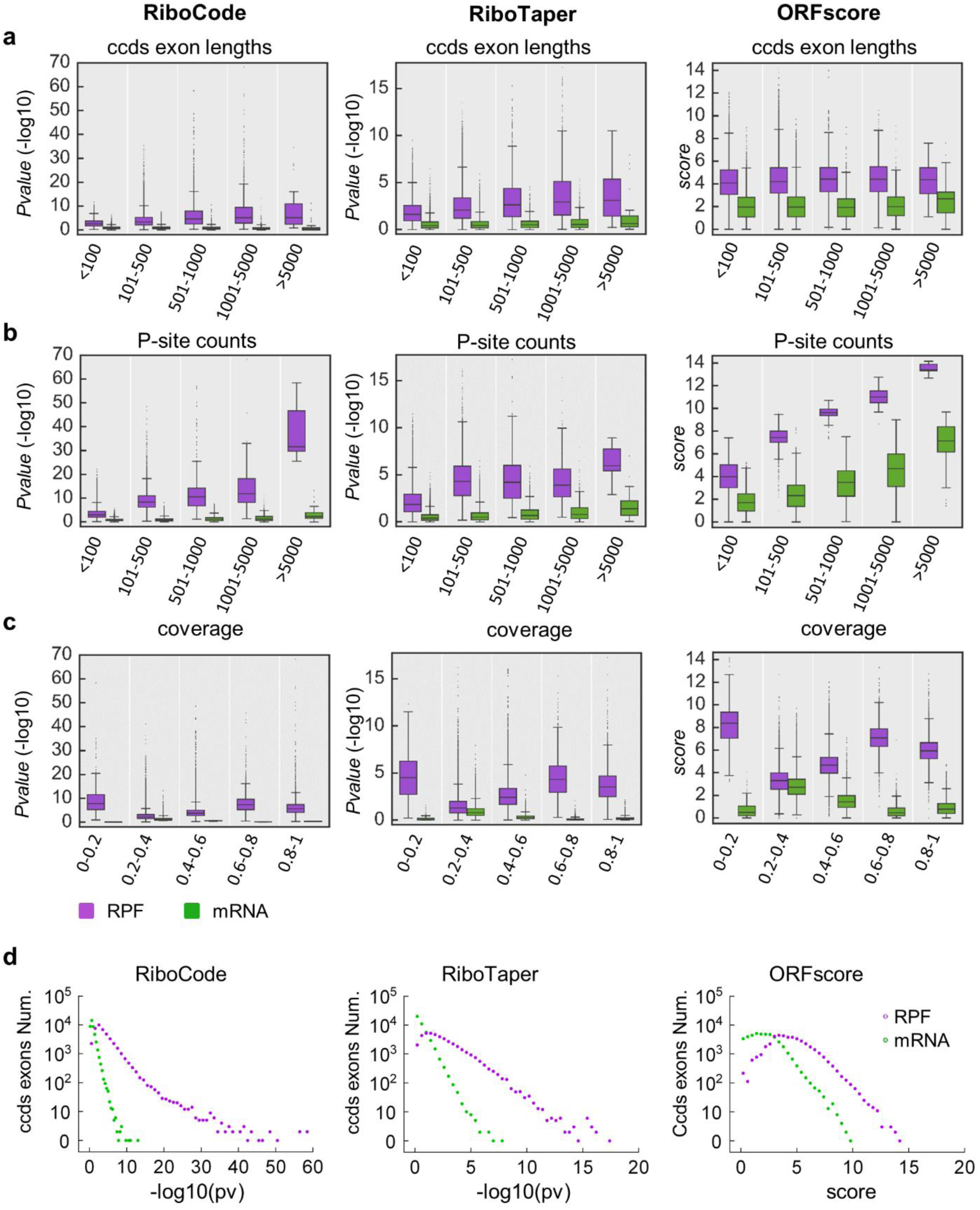
Distributions of the quantitative evaluations of translation with different methods. The CCDS exons were binned into 5 subgroups according to their length (a), coverage (b), or total read count (c). Within each bin, box plots were then generated to illustrate the distributions of the quantitative assessments of translation obtained with different methods (-log10 of the P-value for RiboCode and RiboTaper, and score for ORFscore) with the RPF data or RNA-seq data. Finally, the overall distributions of the quantitative assessments of translation obtained with different methods are provided (d).

**Supplementary Figure 2.**
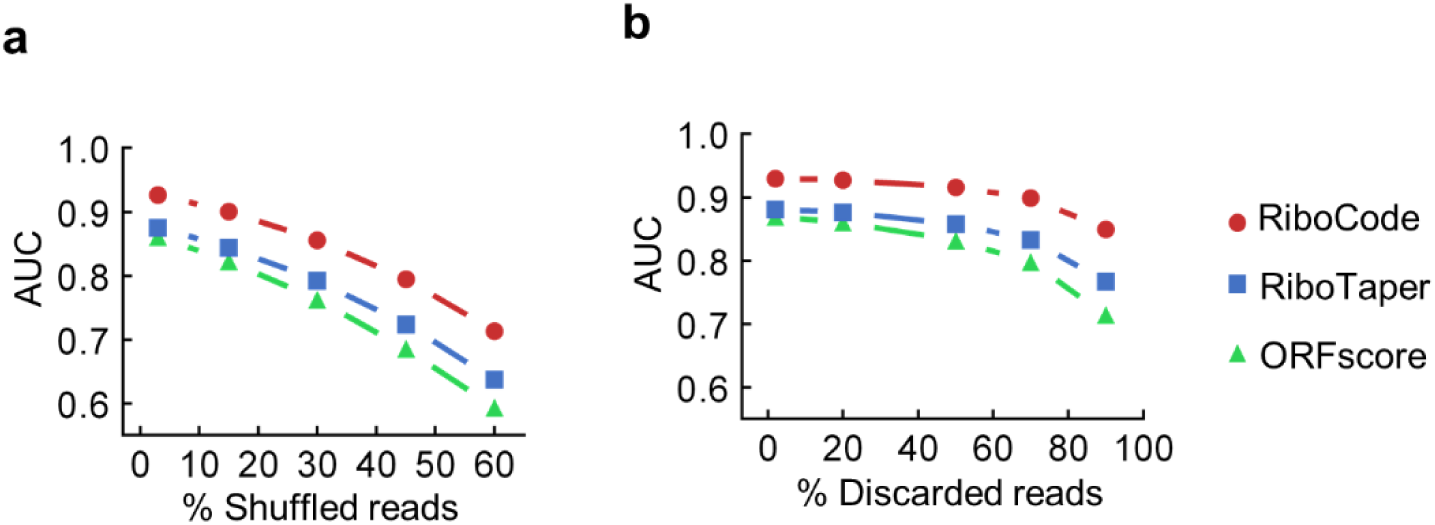
Performances of different methods on simulation data with artificial noise or reduced sequencing depth. (a, b) The area under the curve (AUC) of the ROC curves generated with the results of RiboCode, RiboTaper and ORFscore, which were applied on simulation datasets with different levels of noise (a) or with reduced number of the RPF reads (b).

**Supplementary Figure S3.**
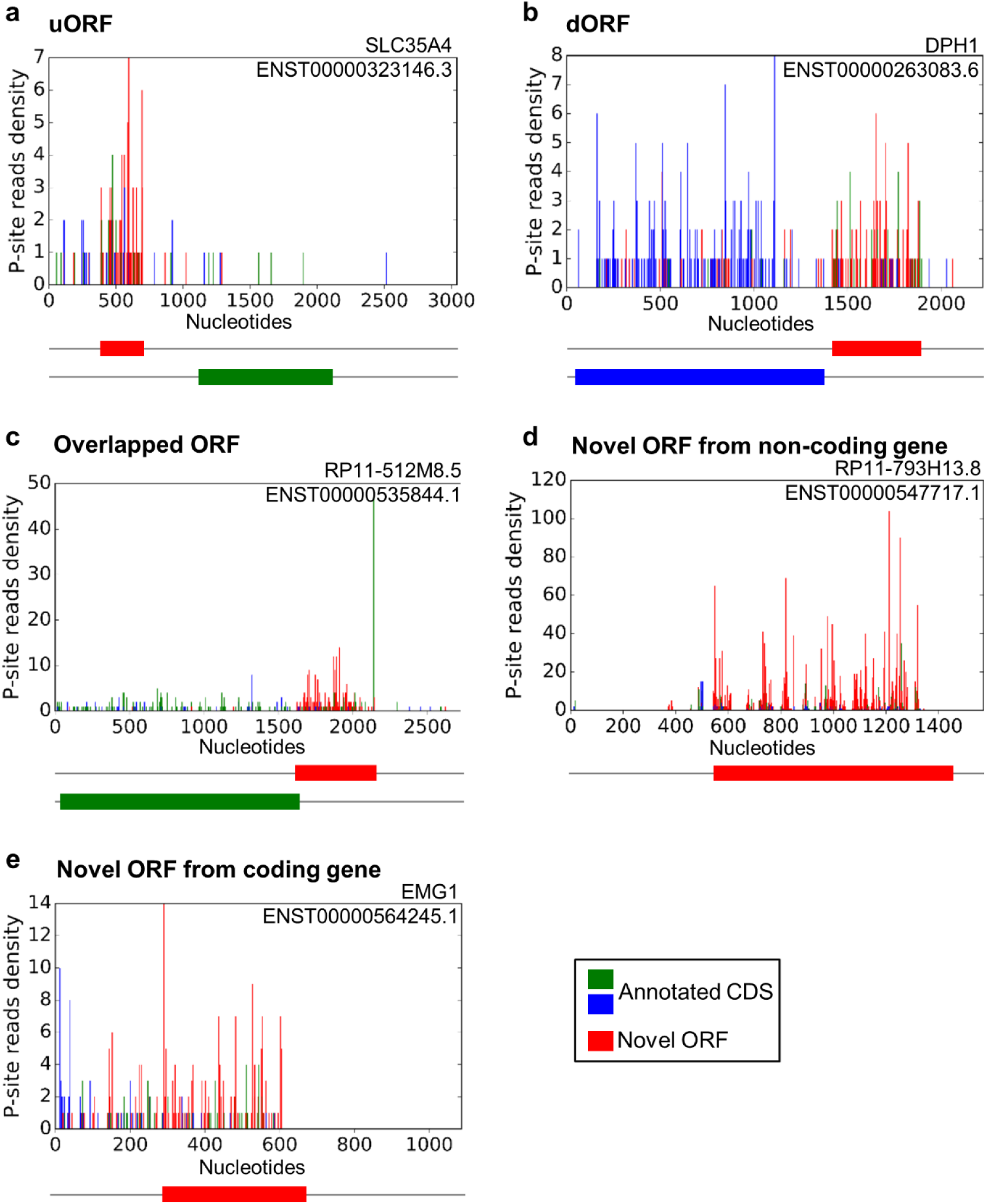
Examples of uORF, dORF and off-frame overlapping ORFs annotated by RiboCode. Examples of the non-canonical ORFs (uORFs, dORFs, overlapping ORFs, and the novel ORFs from the coding and non-coding genes) that were annotated by RiboCode with the HEK293 data.

**Supplementary Figure 4.**
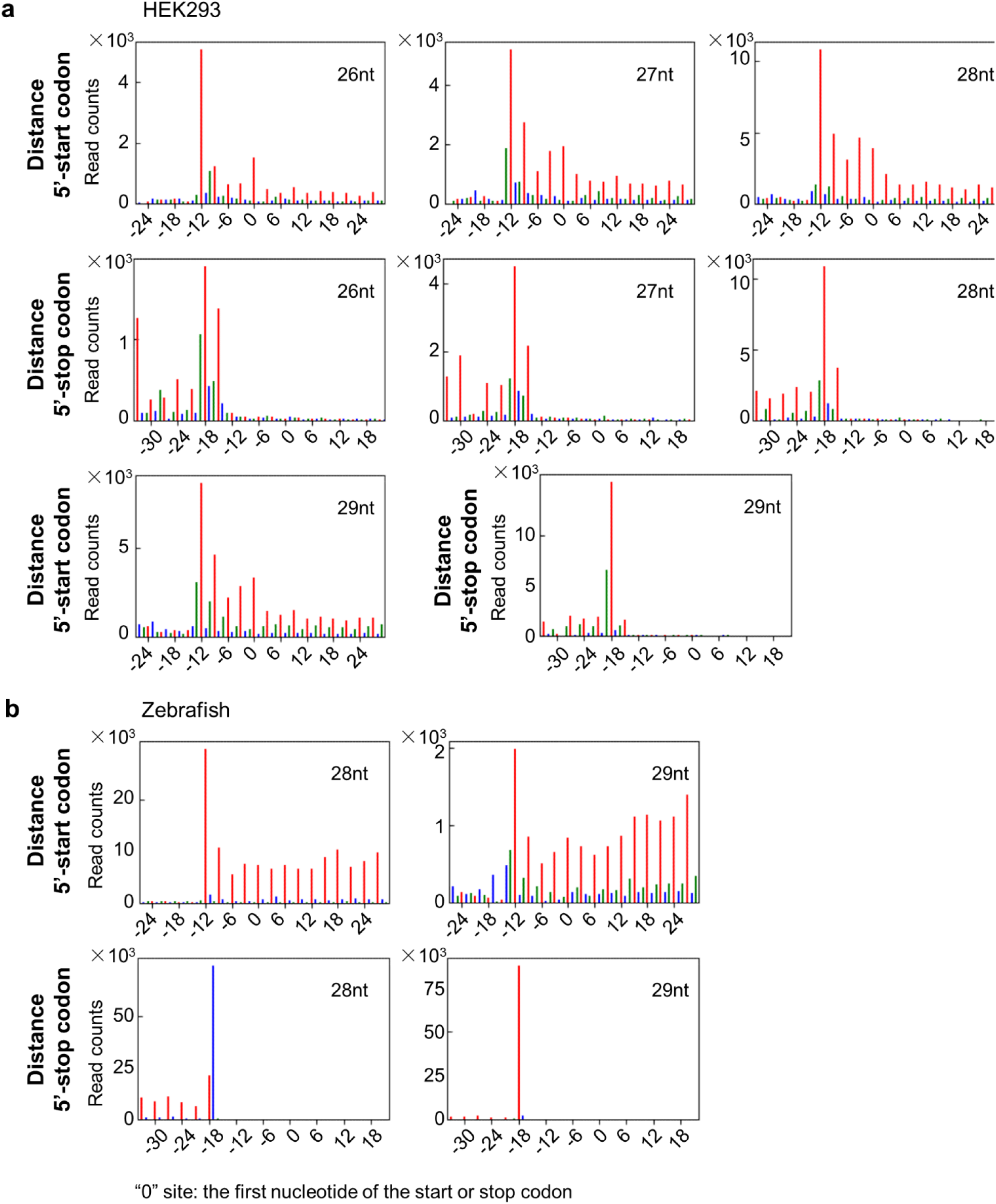
Metagene analysis for RPF length selection and P-site identification. Aggregate plots of distance from the 5’ end of RPF reads to the annotated start or stop codons, for different lengths of RPFs from the ribosome profiling data of HEK293 (a) and Zebrafish (b).

**Supplementary Figure 5.**
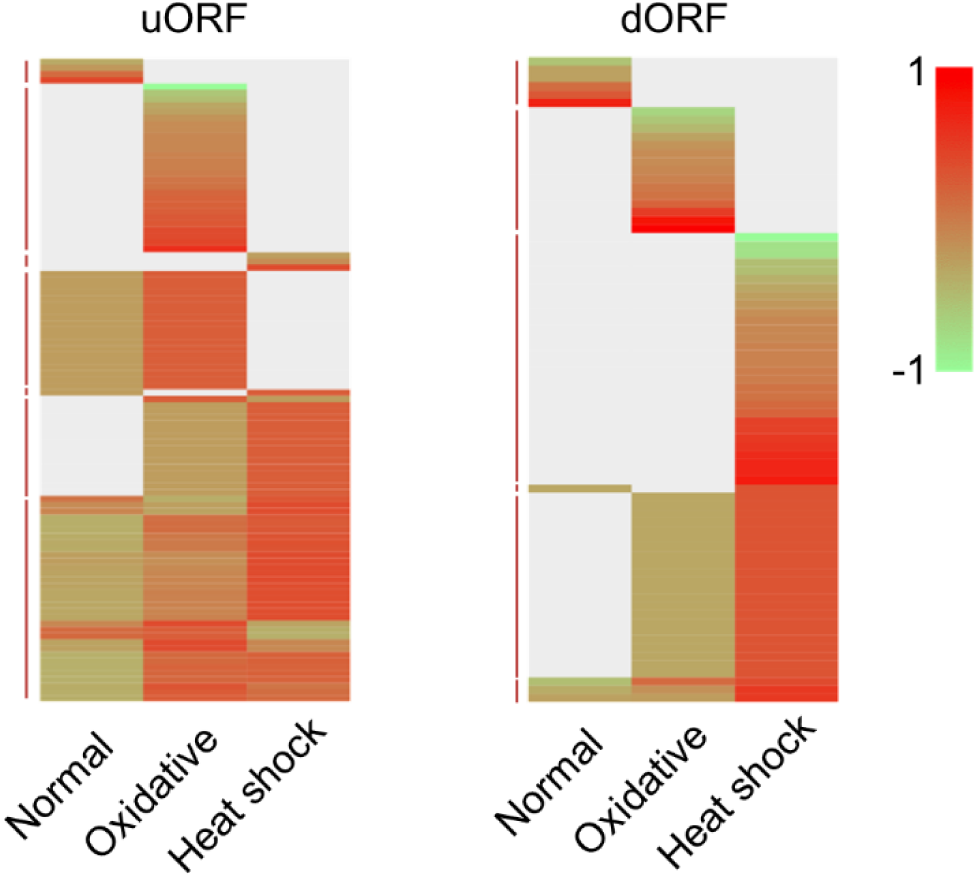
Relative levels of the RPF read counts on the uORFs and dORFs. To show the relative levels across different conditions, the normalized RPF read counts on each uORF or dORF were further centralized across the 3 conditions: normal, oxidative stress, and heat shock. The color was left blank for the uORF or dORF that was not annotated by RiboCode under a specific condition.

### Supplementary Tables

As advised by the editorial office, the following Supplementary Tables will be provided later for the full peer review.

Supplementary Table 1. CCDS exons identified by three different methods. Supplementary Table 2. Complete annotation of the ORFs by RiboCode and RiboTaper with the HEK293 data.

Supplementary Table 3. Complete annotation of the ORFs by RiboCode and RiboTaper with the Zebrafish data.

Supplementary Table 4. ORFs annotated by RiboCode and RiboTaper that were supported by MS data.

Supplementary Table 5. Complete annotation of the ORFs by RiboCode with the yeast data under 3 conditions.

Supplementary Table 6. RPF read counts of the ORFs annotated by RiboCode with the yeast data under 3 conditions.

Supplementary Table 7. Basic statistics, RPF read lengths, and P-site positions of the datasets used in the current study.

